# Nanog fluctuations in ES cells highlight the problem of measurement in cell biology

**DOI:** 10.1101/060558

**Authors:** Rosanna C G Smith, Patrick S Stumpf, Sonya J Ridden, Aaron Sim, Sarah Filippi, Heather Harrington, Ben D MacArthur

## Abstract

A number of important pluripotency regulators, including the transcription factor Nanog, are observed to fluctuate stochastically in individual embryonic stem (ES) cells. By transiently priming cells for commitment to different lineages, these fluctuations are thought to be important to the maintenance of, and exit from, pluripotency. However, since temporal changes in intracellular protein abundances cannot be measured directly in live cells, these fluctuations are typically assessed using genetically engineered reporter cell lines that produce a fluorescent signal as a proxy for protein expression. Here, using a combination of mathematical modeling and experiment, we show that there are unforeseen ways in which widely used reporter strategies can systemically disturb the dynamics they are intended to monitor, sometimes giving profoundly misleading results. In the case of Nanog we show how genetic reporters can compromise the behavior of important pluripotency-sustaining positive feedback loops, and induce a bifurcation in the underlying dynamics that gives rise to heterogeneous Nanog expression patterns in reporter cell lines that are not representative of the wild-type. These findings help explain the range of published observations of Nanog variability and highlight a fundamental measurement problem in cell biology.

---

Fluorescence has been used to report expression of gene products in live cells since green fluorescent protein (GFP) was first cloned and utilized as a tracer [1, 2]. Live cell fluorescence imaging and analysis techniques allow investigation of temporal changes in protein expression and have consequently become an essential tool in modern molecular biology [3]. However, their proper use requires the reporter signal to be representative of expression of the protein of interest at the scale of interest. In particular, if the reporter is to be used as a proxy for protein expression within a single cell then in order to be able to draw accurate conclusions the reporter signal should be representative of protein expression in *that* particular cell. This issue is particularly relevant when functional assays are performed after cell sorting based upon reporter signal intensity and can present a significant problem if the long-term outcome of any subsequent assays are driven by rare subpopulations of misidentified cells. As interest in single cell biology has increased, some generalized concerns about the fidelity of standard live-cell reporter strategies have been raised [4, 5]. However, the ways in which the genetic manipulations involved in generating reporter cell lines can affect endogenous gene expression kinetics are not well understood.

Here, we show that there are unforeseen ways in which commonly used live cell reporter strategies can perturb the regulatory mechanisms they are intended to measure and induce qualitative changes in dynamics that are not representative of the wild-type. Counter-intuitively, attempts to improve reporter signal can exacerbate this issue and result in further loss of resolution on the true dynamics. Since predicting when these problems will occur requires a priori knowledge of the underlying regulatory control mechanisms of the system under study – which is typically the knowledge that the reporter was introduced to provide – our results highlight a basic measurement problem in cell biology, reminiscent of that encountered in quantum physics [6], in which the act of measuring disturbs the system being measured.

In the first part of the paper we use a mathematical argument to show why it should not generally be expected that reporters will faithfully reflect gene expression dynamics at the single cell level, and why reporter accuracy depends strongly upon regulatory context. Surprisingly, this analysis also demonstrates that expression noise can improve, rather than degrade, reporter accuracy. To illustrate these general results we then consider the case of Nanog, a central pluripotency regulator in ES cells, which has been seen to fluctuate stochastically within individual cells, yet for which different types of reporter have given different assessments of the strength and developmental significance of these fluctuations [4, 7–10]. Using both mathematical modeling and experiment, we show how different reporter strategies can disturb the kinetics of endogenous Nanog regulatory mechanisms in different ways, giving different assessments of Nanog dynamics.

## RESULTS

### Reporter accuracy depends upon regulatory context

The basic type of targeted gene reporter is a knock-in reporter, in which one of the alleles for the gene of interest is replaced with a reporter gene, often encoding for a fluorescent protein, perhaps with additional features such as an antibiotic selection cassette [3]. Due to the loss of one gene copy, these are often described as heterozygous loss-of-function reporters. For such constructs to be effective at the single cell level, the fluorescence signal driven from the reporter allele should accurately represent protein expression from the wild-type allele. We therefore begin by considering a toy model of transcriptional co-activity, in order to explore the conditions under which two alleles that are subject to the same regulatory control may either synchronize or decouple in their activity, and thereby the conditions under which the output of one allele may be used to report on the other. For simplicity we will focus on mRNA dynamics, but similar reasoning may also be extended to the protein level. Consider the transcriptional dynamics of two alleles of the same gene in a single cell. Let *m*_1_ denote the number of mRNA transcripts associated with allele 1, let *m*_2_ denote the number of mRNA transcripts associated with allele 2, and assume that expression from both alleles is governed by linear birth-death processes with production rates 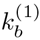, 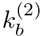 and decay rates 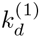, 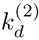. Thus, we are concerned with the dynamics of the following system of reactions,

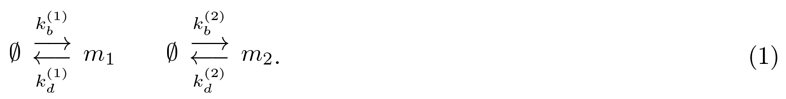

**Figure 1.**
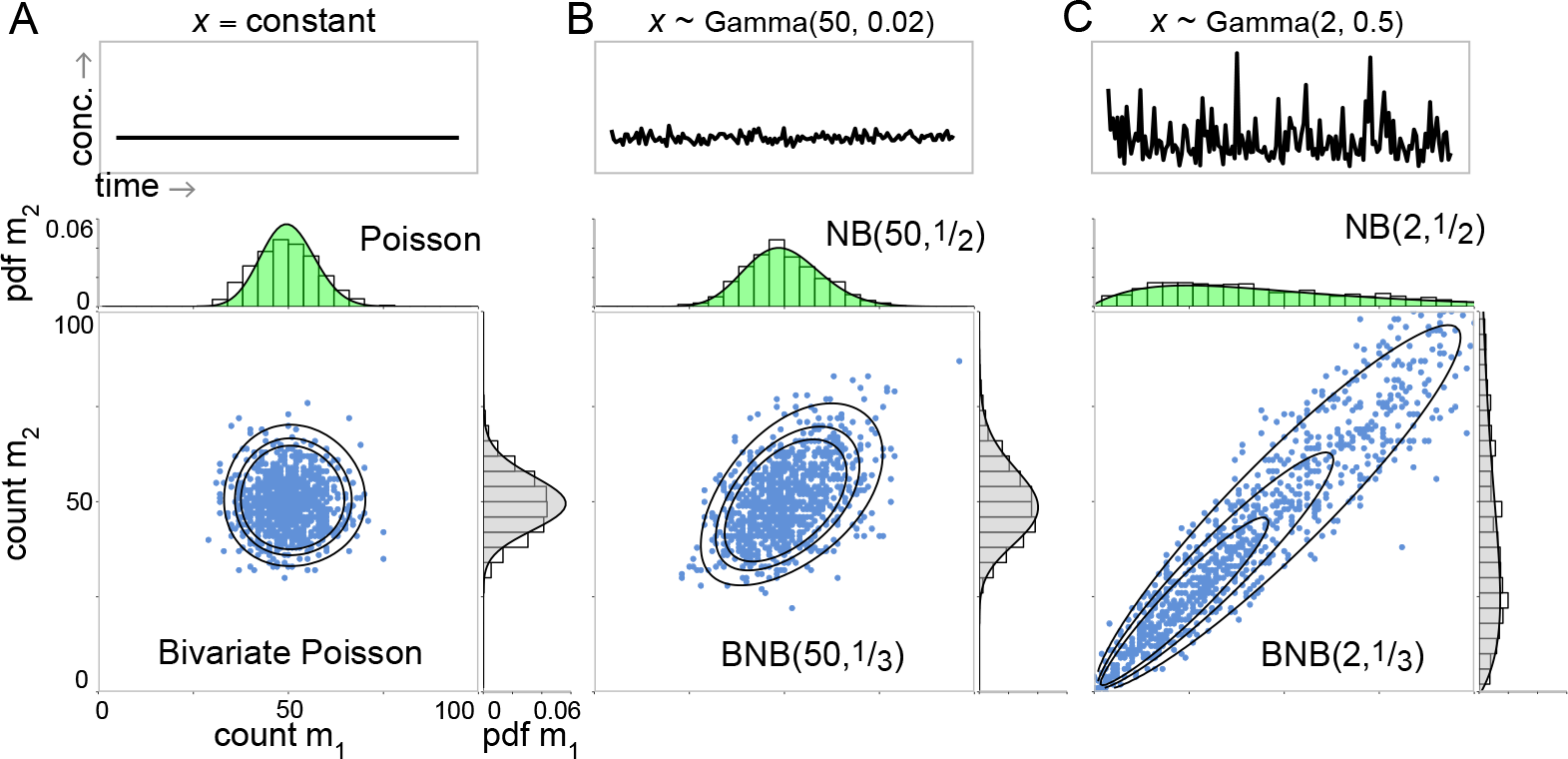
Reporter accuracy depends upon regulatory context. Identical alleles of the same gene produce mRNA molecules *m*_1_ and *m*_2_. Alleles 1 and 2 behave independently when there is no common upstream regulator or regulator concentration (*x*) is constant. **Top panels:** Fluctuations of upstream regulator concentration. Examples shown for constant *x* (**A**) and *x* ~ Gamma(*r*, *θ*), for low regulator dispersion *θ* = 0.02 (**B**) and high regulator dispersion *θ* = 0.5 (**C**). **Bottom panels:** Joint and marginal distributions for *m*_1_ and *m*_2_. All distributions use λ = 50 for both alleles. Contours show probabilities: 0.0001 inner, 0.0003 middle, 0.0005 outer. BNB(*r*,*p*) denotes Bivariate Negative Binomial, given by Eq. 4 with *p* = *q* for identical alleles. NB(*r*,*p*/(1 − *p*)) denotes Negative Binomial. Scatter plots and histograms are shown for a random sample of 1000 draws in each case. The same scales apply to all comparable plots.

This is clearly a simplistic view of transcription (others have extended this basic model, including descriptions of transcriptional bursting [11]), yet it suffices to illustrate some of the essential issues regarding the reliability of reporters and is analytically tractable. Since the alleles are not coupled together they act independently and the stationary joint probability mass function (PMF) for this process is the product of two Poisson distributions (Fig.1A),

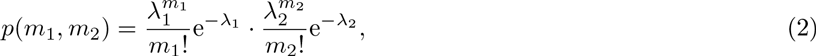

where 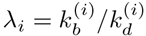 for *i* ∈ {1, 2}. As we have not allowed for co-regulation of expression this model is rather artificial. In reality, we expect that if the alleles are both under the same promoter control then they will be regulated by the same upstream factors, and this co-regulation may coordinate their dynamics. In order to couple the alleles together we allow the transcription rates to be driven by a shared upstream regulator *X*. Let *x* denote the concentration of *X* and let *ρ*(*x*) be the stationary probability density function for *x*. Assuming that the mRNA birth rate is now given by 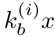, the stationary joint PMF is then obtained from Bayes’ theorem,

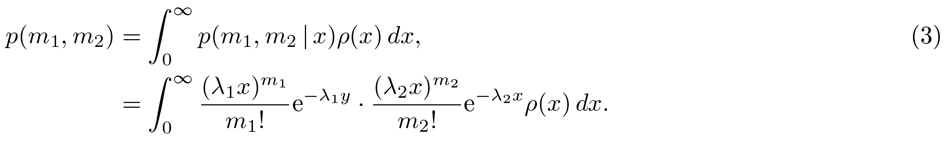

Since the joint PMF *p*(*m*_1_, *m*_2_) depends upon the distribution of the upstream regulator, an appropriate form for *ρ*(*x*) must be chosen. It is commonly observed that protein concentrations are Gamma distributed, so this is a natural choice [11]. In the case *x* ~ Gamma(*r*, *θ*) the joint PMF is

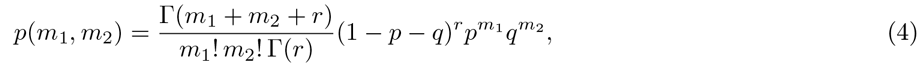

where *p* = λ_1_*θ*/[1 + *θ*(λ_1_ + λ_2_) and *q* = *ap* with *a* = λ_2_/λ_1_. The marginal distributions for the two allele products are then negative binomials and the joint PMF is a bivariate negative binomial distribution (BNB), as shown in Fig.1B-C. If the two alleles are kinetically identical (λ_1_ = λ_2_), then the marginal distributions will be identical, and the product of either allele may be used to report on the other at the *population* level (assuming that the same dynamics occur within each cell in the population). However, this does not guarantee any association between the allelic outputs at the *individual* cell level. To measure the degree of association between alleles within an individual cell we must consider the covariance between their outputs, which is easily calculated in this case (see SI Text) and has a particularly simple form,

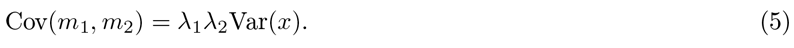

Thus, the covariance between the two allele products is proportional to the variance of the upstream regulator and the sensitivities of the two alleles to the upstream regulator. While the form of joint PMF given in Eq. 4 depends upon the upstream regulator being Gamma distributed, Eq. 5 holds for any upstream distribution *ρ*(*x*) (see SI Text). A comparable result may be obtained when transcription from each allele occurs at rate 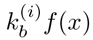, for any smooth function *f*(*x*) (see SI Text). Similarly, it may also be shown that the correlation between the two alleles depends in a monotonic positive way on the the Fano factor (or index of dispersion) of the upstream regulator (see SI Text). These results highlight two points of importance to the design and proper use of reporters: (1) regulatory noise upstream can increase the coordination of alleles downstream and therefore *improve* reporter accuracy; (2) a reporter’s accuracy depends on the regulatory context in which it is placed, here represented by the dynamics of the upstream regulator *X*. Thus, the accuracy of a reporter depends upon factors that are external to its design. In a complex regulatory environment, these external factors may not be fully (or even partially) known, and their effect on the reporter may be correspondingly hard to predict or control.

Taken together this analysis suggests that reporter efficacy is highly contextual. In order to explore this phenomenon further it is helpful to consider a specific example. Here we examine the case of Nanog, an important, highly-studied, yet still poorly understood, regulator of pluripotency in ES cells.

### Dynamics of Nanog and reporter co-expression

It has been widely observed that Nanog expression appears to fluctuate stochastically in individual ES cells [7, 12, 13]. However, different reporter constructs have given different assessments of the strength and developmental significance of these fluctuations [4, 7–10, 14–16] and some concerns have been raised that the use of reporters may be introducing artifacts that are confounding, rather than clarifying, our understanding of pluripotency [4, 5].

To address this issue we will consider a simple mathematical model of Nanog dynamics in ES cells in the presence of different kinds of reporter constructs. Nanog levels are regulated in pluripotent cells by a complex network of molecular interactions that involve both protein-protein and protein-DNA interactions [17–20]. A number of groups have modeled these dynamics mathematically [12, 21, 22]. At the core of this extended regulatory network is a series of nonlinear positive feedback loops that are dependent on Nanog for their function [23, 24]. Since these feedback loops are central to Nanog regulation, and in order to maintain a tractable mathematical model of general relevance, we will focus on this aspect of Nanog regulation here. Since positive feedback mechanisms naturally give rise to switch-like dynamics, they are correspondingly central to many kinds of cell fate decisions [25–27]. Therefore, the model of positive feedback that we outline below is of general relevance to the design of reporters for other similarly regulated lineage specifying master transcription factors.

We consider the following set of ordinary differential equations (ODEs) as a simple model of Nanog protein dynamics in wild-type cells,

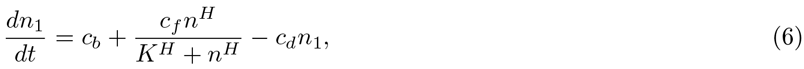

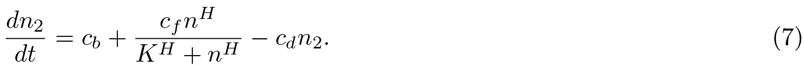

where *n*_*i*_ denotes the concentration of the Nanog protein output of allele *i* ∈ {1, 2}. The first terms on the right hand sides of these equations account for baseline production at a constant rate *c_b_*; the second terms account for feedback-enhanced production at a rate dependent on total Nanog concentration *n* = *n*_1_ + *n*_2_, up to a maximum rate *c*_*f*_; and the third terms account for Nanog protein decay at rate *c*_*d*_. Adding these equations we obtain an ODE for the total Nanog protein concentration *n*,

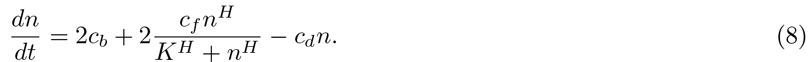

In order to better understand the model dynamics it is convenient to nondimensionalize this equation. Doing so, using the scalings 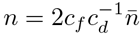, and 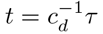 we obtain,

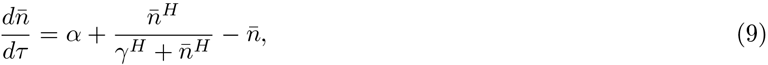

where 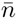 is the dimensionless total Nanog concentration and *τ* is dimensionless time. The dimensionless constants *α* = *c*_*b*_/*c*_*f*_ and *γ* = *γ*_wt_ = *c*_*d*_*K*/2*c*_*f*_ describe the relative strength of the basal and positive feedback enhanced production rates respectively. Eq. 9 has either 1 or 2 stable equilibrium solutions depending on the relative sizes of *α*, *γ* and the Hill coefficient *H*. In particular, for *H* > 1, two bifurcation curves in the *αγ* plane may be found (see SI Text). The case *H* = 2 suffices to illustrate the general structure of the resulting classification diagram (see Fig. 2A). In this case, the bifurcation curves are,

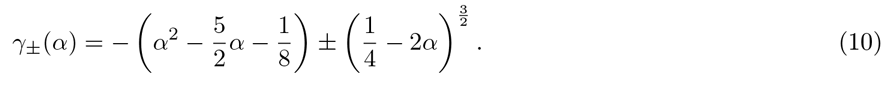

**Figure 2.**
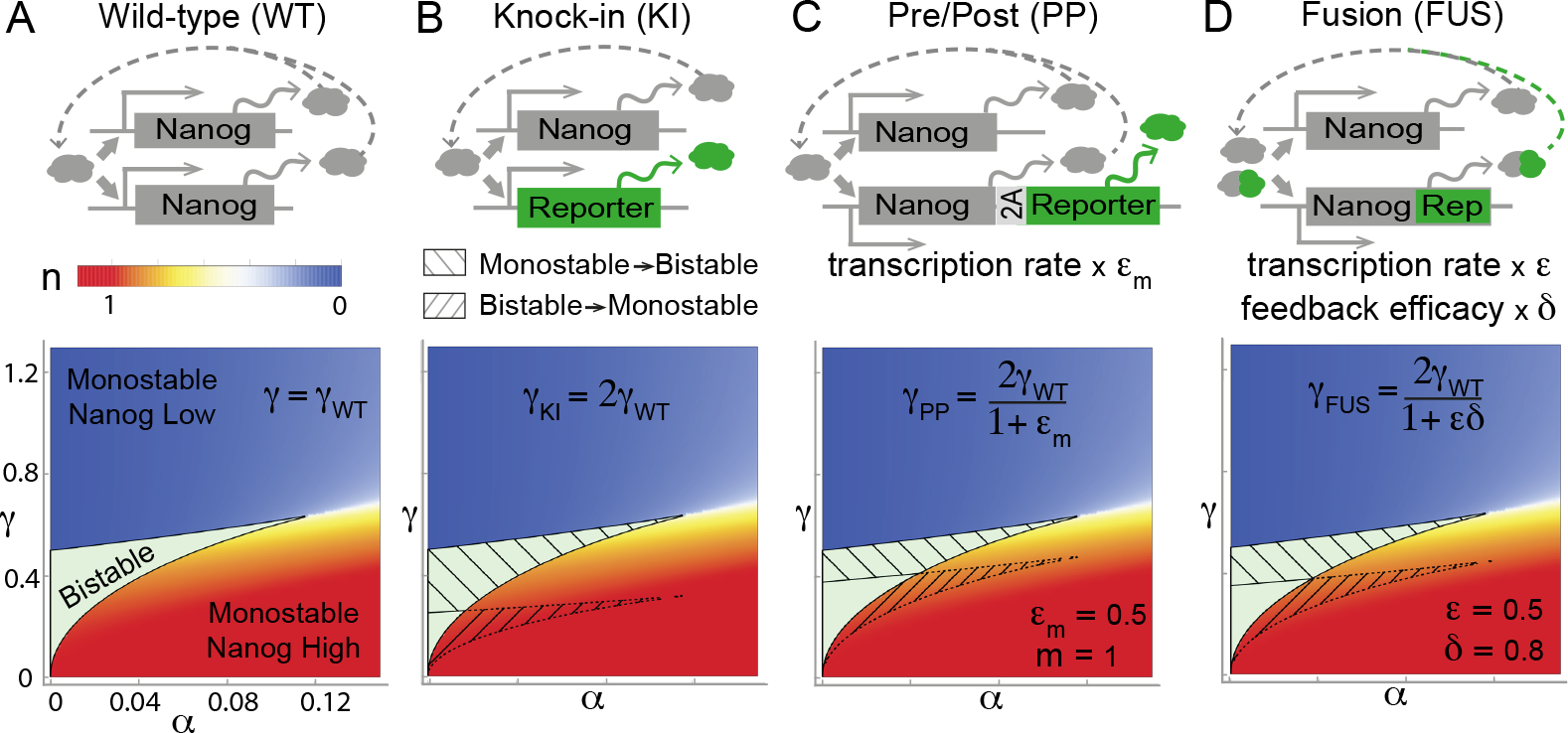
Perturbation of Nanog dynamics by reporters. **A.** *Wild-type*: Nanog protein is produced from both alleles. Monostable or bistable dynamics can occur depending on *α* and *γ*. **B**. *Knock-in reporters*: one allele is left intact and one allele produces an inert reporter protein. Nanog production is reduced by a factor of two, thereby doubling *γ*. **C**. *Pre/post reporters*: both alleles encode for Nanog; *m* copies of a self-cleaving reporter protein are also transcribed from one allele. The rate of transcription from the reporter allele is reduced by factor 0 ≤ *ϵ*_*m*_ ≤ 1 thus increasing *γ* by a factor 2/(1 + *ϵ*_*m*_). As *ϵ*_*m*_ decreases with *m*, these reporters become more prone to systemic errors with each additional insert. **D**. *Fusion reporters*: one allele encodes for Nanog; the other for a fusion of Nanog and a reporter protein. The rate of transcription from the reporter allele is decreased by factor 0 ≤ *ϵ* ≤ 1 and the reporter fusion reduces Nanog feedback functionality by a factor 0 ≤ *δ* ≤ 1 thus increasing *γ* by a factor 2/(1 + *ϵδ*). Hatching shows *at risk* regions.

If the model parameters fall *inside* the region enclosed by these curves then Nanog expression dynamics are bistable; if the model parameters fall *outside* this region then Nanog expression dynamics are monostable. In the presence of molecular noise, which is inherent to the intracellular micro-environment, bistability can give rise to coexisting subpopulations of phenotypically distinct cells within an isogenic population under the same environmental conditions [26]. Thus, both homogeneous and hetereogeneous Nanog expression patterns are allowed by this model, depending on whether the underlying dynamics are monostable or bistable. It should therefore be expected that Nanog expression patterns in ES cell populations will vary substantially under different experimental conditions, depending on how they stimulate Nanog feedback mechanisms, as is commonly observed [4, 8, 28]. More significantly, it should also be expected that any genetic interventions that perturb the kinetics of Nanog feedback have the potential to push the dynamics in or out of the bistable regime, thereby affecting a *qualitative* change in expression patterns. To see this, consider the case of a heterozygous knock-in reporter, in which one allele produces an inert reporter and one allele is left intact. In this case, the wild-type kinetics described by Eqs. 6 are modified to,

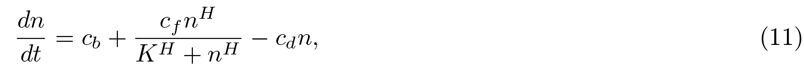

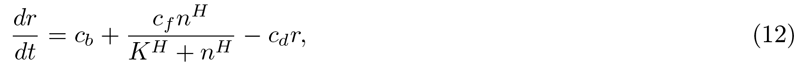

where *r* is the reporter protein concentration. For simplicity we have assumed that the reporter and Nanog protein half-lives are perfectly matched (this assumption may be relaxed without altering conclusions). The dimensionless equation for total Nanog concentration in the reporter line is,

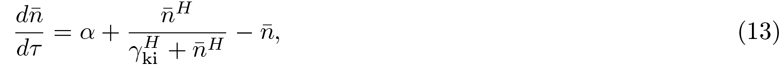

where *α* = *c*_*b*_/*c*_*f*_ as before, but *γ*_ki_ = *c*_*d*_*K*/*c*_*f*_ = 2*γ*_wt_ (see SI Text for details). In this case, the loss of Nanog production from one allele diminishes the Nanog production rate by a factor of two, which weakens the endogenous feedback mechanisms and thereby doubles the parameter *γ*. Since for fixed *α* the magnitude of *γ* determines if the dynamics are monostable or bistable, and therefore if Nanog is homogeneously or heterogeneously expressed in the population, this change can induce a heterogeneous Nanog expression pattern in the reporter cell line that is not found in the wild-type (or vice versa). Areas in the *αγ* plane for which the map (*α*, *γ*) ↦ (*α*, 2*γ*) crosses one of the bifurcation curves in Eq. 10 are *at risk* of this kind of perturbation (hatched regions in Fig. 2). Importantly, this problem is not restricted to knock-in lines: similar issues arise with a wide range of other reporters, including protein fusions and reporters inserted before or after the Nanog gene (using 2A self-cleaving peptides or internal ribosomal entry sites, for example), in both single allele and dual allele reporter systems. Fig. 2 summarizes similar analyses for some other reporters (see SI Text for details of calculations for these and a range other reporters, including dual allele systems).

In order to determine the extent to which these issues arise in experiment, we compared Nanog expression patterns in wild-type (male v6.5) mouse ES cells to those in a heterozygous knock-in reporter ES cell line with the same male v6.5 genetic background, in which the Nanog coding sequence was replaced with a GFP-IRES-puro reporter on one allele [29] (designated NHET cells). Cells were cultured in standard culture conditions (0i, serum plus LIF) and 2i conditions (0i conditions with the addition of mitogen-activated protein kinase and glycogen synthase kinase 3 inhibitors), which maintain ‘ground state’ pluripotency [30]. Homogeneous Oct3/4 expression was confirmed via immunostaining in all culture conditions (Fig. S1). Nanog expression in individual cells was assessed via fluorescence immunolabeling and quantified by flow cytometry (Fig. 3) and image analysis (Fig. S2). Substantial variability in Nanog expression was observed in both v6.5 and NHET lines in 0i conditions (Fig. 3A and Fig. S1). In accordance with previous reports, substantially less variability was observed in 2i conditions (Fig. 3A and Fig. S1) [30]. Distinctly bimodal GFP fluorescence was observed for NHET reporter cells in 0i cultures, with cell clusters containing both GFP high cells and GFP low cells present in abundance. In both conditions a clear mismatch between Nanog and GFP expression levels was observed in a substantial proportion of cells (Fig. 3B). This was most apparent in 0i conditions, where of the highest 20% of Nanog expressing cells, 23% were GFP low, while of the lowest 20% of Nanog expressing cells 11% were GFP high. In 2i conditions the percentage of GFP low cells was consistent across the Nanog distribution, indicating that GFP status was not indicative of Nanog expression. In addition, within the GFP high subset the there was no clear association between Nanog and GFP expression levels (Fig. 3B).

**Figure 3.**
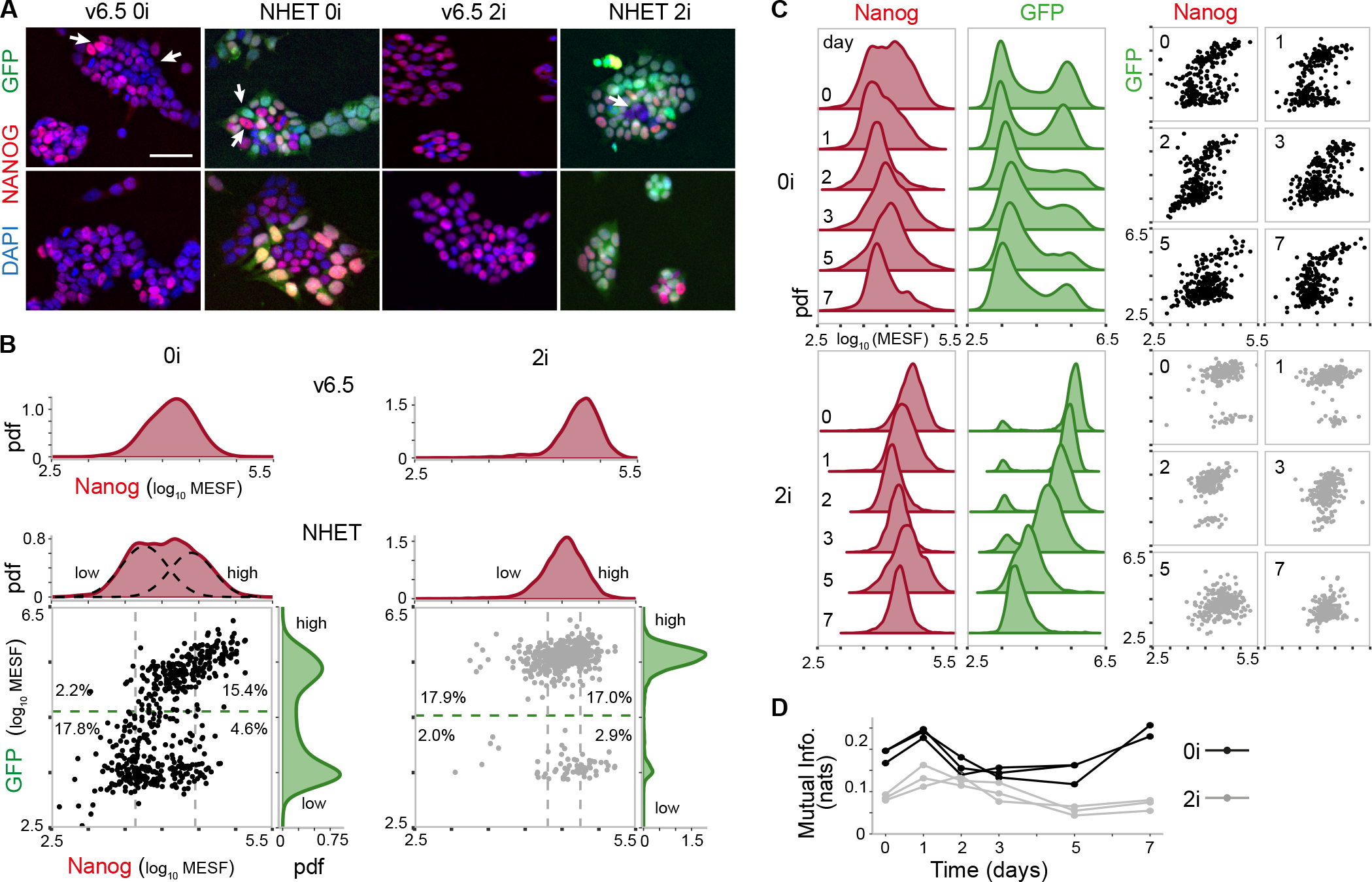
Nanog expression in wild-type and reporter cell lines. **A.** Wild-type and Nanog reporter (NHET) cell cultures in 0i and 2i conditions. Nanog immunofluorescence is in red, and direct GFP fluorescence is in green. White arrows indicate Nanog low/high cells (wild-type) or cells in which there is a Nanog-GFP mismatch (NHET). Grayscale fluorescence signals are shown Fig. S1. Scale bar represents 50*μ*m. **B**. Representative flow cytometry distributions of Nanog in wild-type cells (top) and Nanog-GFP joint distributions in NHET cells (bottom). Dashed black line shows components of fit of a 2 component Gaussian mixture model to the wild-type Nanog distribution in 0i conditions. Dashed threshold lines indicate regions of GFP high/low expression and Nanog high/low expression (highest 20% and lowest 20% of cells). Percentages show proportions of total cells in the relevant subpopulations. **C**. Changing Nanog and GFP distributions during undirected differentiation subsequent to LIF withdrawal starting from 0i (top) and 2i (bottom) cultures. **D**. Mutual information between GFP and Nanog during differentiation. Results of three experimental repeats are shown.

To determine whether Nanog expression was perturbed by introduction of the knock-in reporter we compared Nanog distributions between NHET and wild-type v6.5 cell lines using immunostaining and flow cytometry. In NHET cells, the Nanog distribution in 0i conditions exhibited a wide flattened distribution. Fitting of this data to a Gaussian mixture model (GMM) with one or two components (see SI Text), revealed that the 2-component model best described the data, suggesting the presence of two coexisting subpopulations of cells characteristic of bistability in the underlying dynamics (Fig. 3B). By contrast, in wild-type v6.5 cells the Nanog distribution in 0i conditions was less broad and was better fit by a single component model, suggesting monostability in the underlying dynamics (Fig. 3B). To establish the robustness of these results we also assessed Nanog expression using image analysis, and these broad conclusions were confirmed (Fig. S2A). Taken together these analyses suggest that the bimodal expression patterns observed in 0i conditions using the NHET line are due to systemic perturbation of the Nanog regulatory network by the reporter, as predicted by theory. By contrast, both wild-type and NHET cells expressed similar, more compact, Nanog distributions in 2i conditions, with neither showing evidence of bimodality. This suggests that 0i conditions only weakly stimulate the endogenous regulatory feedback mechanisms, causing the system to lie within the *at risk* region of the *αγ* parameter plane, while in 2i conditions these feedback mechanisms are strongly activated, causing the system to lie outside the *at risk* region.

To further investigate the extent to which environmental changes might affect the fidelity of the reporter output we also sought to assess the association between Nanog and GFP during the process of cellular differentiation. Starting in 0i or 2i conditions, NHET cell cultures were allowed to undergo undirected differentiation by withdrawing LIF for a period of 7 days. Fig. 3C and Fig. S2B show the evolving joint distributions for Nanog and GFP co-expression. From 0i conditions, the proportion of cells in the GFP high population gradually decreased over time and the corresponding Nanog distribution concomitantly evolved from an initial broad, flat distribution to a narrow distribution with lower average expression, indicating gradual loss of Nanog expression. From 2i conditions both GFP and Nanog levels gradually declined over time without qualitative change in distribution shape. To quantify the association between Nanog and GFP levels we calculated the mutual information between their expression patterns,

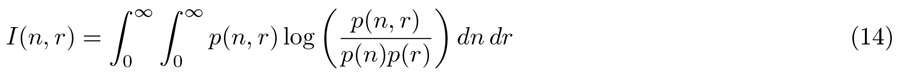

where *p*(*n*, *r*) is the joint probability density function for Nanog and GFP co-expression, and *p*(*n*) and *p*(*r*) are the marginal probability density functions for Nanog and GFP expression respectively. Mutual information is a powerful generalization of traditional measures of association, such as correlation, which is able to identify nonlinear relationships between variables [31]. In this context, the mutual information provides an unbiased measure of the the amount of information that knowledge of a cell’s GFP status provides about its Nanog status (zero mutual information indicates complete independence, low values indicate near independence, high values indicate strong association). In all cases, the mutual information exhibited a mild transient increase, indicating a slight increase in strength of association during the early stages of differentiation (Fig. 3D and Fig. S2C). However, mutual information was always low, indicating that Nanog and GFP signals are only weakly related both in and out of equilibrium (Fig. 3C).

## DISCUSSION

The advantages of genetic reporters are substantial: they provide a means to investigate expression dynamics of hard-to-monitor proteins and enable live cell observation, tracking, and selection. By assessing expression directly via fluorescence rather than indirectly via immunolabeling they also provide a more transparent way to assess protein activity, free of the reproducibility issues associated with the use of antibodies. However, it is generally accepted that genetic reporter systems are not perfect: quantification is normally relative, reporter fluorescence is an imperfect proxy-measurement for the variable of real interest, and it is known that wild-type dynamics may be compromised, for example by fusing cumbersome fluorescent proteins to (often relatively small) proteins of interest [32]. To assess the importance of these issues the advantages and disadvantages of different types of reporter are usually considered purely in terms of their technical characteristics, or with only limited concern for their regulatory context, for instance to match reporter half-life to that of the protein of interest [33]. However, the systemic limitations of reporter strategies, due to the interaction of reporter constructs with specific endogenous regulatory mechanisms, have not been well appreciated. Yet, our results show that if such limitations are not taken into account, then misleading results can follow.

Taken together this work suggests several practical guidelines to help prevent unforeseen issues with reporter observations: firstly, the scale at which the reporter is used should be considered. In particular, for assays involving cell sorting based upon reporter signal, accuracy should be tested at the single cell level prior to subsequent functional assays. Secondly, systemic limitations of reporters, due to interactions between the reporter and its regulatory context, should also be considered. The most appropriate reporter strategy will be determined by a trade-off between the type of spatial and temporal information required, the strength of the reporter signal required and the likelihood that the reporter chosen will qualitatively perturb the endogenous kinetics. For example, reporters that produce multiple copies of a fluorescent protein per copy of the protein of interest naturally produce a stronger fluorescence signal, yet their construction involves greater genetic intervention, so they also carry a correspondingly higher systemic risk (see Fig. 2 and SI Text). Before designing or using such reporters the benefits of increased signal should therefore be weighed against the increased possibility of systemic errors. For genes that are regulated by positive feedback mechanisms (which includes many developmentally important factors [21, 24, 25, 27]), the risk of systemic failures is greatest for knock-in reporters and least for BAC reporters and, surprisingly, single allele reporters carry less systemic risk than dual allele reporters, which are typically seen to be more accurate (SI Text). Finally, since reporter accuracy depends intimately on regulatory context, and the same reporter in the same cells may fail in some experimental conditions and succeed in others, quality controls should be conducted for all experimental conditions under consideration.

## METHODS

Pluripotent mouse ESCs were cultured in standard 0i and 2i conditions (see Supplementary Information). Nanog reporter line (NHET) was generated by Maherali et al. [29]. For 0i and 2i cultures, 3 technical replicates were assessed for Nanog and GFP distributions by flow cytometry and image analysis at passage number 11 (v6.5s) and passage number 20 (NHETs, also day 0 in time-course experiments). For undirected differentiation time-course experiments, three replicates from each initial condition (0i and 2i) were cultured separately for 7 days after LIF withdrawal on day 0. Cultures were passaged every 2 days and assessed by flow cytometry and image analysis on days 0, 1, 2, 3, 5 and 7. All experimental results in main paper used directly conjugated primary antibodies: Mouse anti-mouse Nanog (Alexa Fluor^®^ 647, 1:200, 560279), and rat anti-histone H3 (pS28) (Alexa Fluor^®^ v450, 560606) to identify mitotic cells. Nanog expression distributions were fitted to a gaussian mixture model with 1 or 2 components using expectation maximization. Model selection was conducted using Bayes Information Criterion (BIC). Mutual information between GFP and Nanog expression levels was estimated using a James-Stein-type shrinkage estimator. Discretization of each variable at each time point was performed separately via the Bayesian Blocks method.

We thank Neil Smyth for the provision of LIF and Jianlong Wang for the NHET cell line. This work was funded by BBSRC Grant No. BB/L000512/

## DETAILED METHODS

### Cell culture

Pluripotent mouse embryonic stem cells (mESC) were cultivated in Dulbecco’s Modified Eagle Medium (DMEM) with 1% Penicillin/Streptomycin, further supplemented with 10% fetal bovine serum, 1× MEM non-essential amino acids, 1× GlutaMAX^™^ and 50 *μM c*_*d*_-Mercaptoethanol. Leukemia inhibitory factor (LIF), was added at a dilution of 1:1000 (produced in house). This is 0i culture medium. For 2i culture medium, 0i medium was supplemented with 1:10000 10mM PD0325901 (Tocris Bioscience, 4197) and 1:3000 10mM CHIR99021 (Reagents Direct, 27-H76). After transfer from 0i media cells were adapted to 2i media over 6 passages. Cells were initially cultured 0.1 % gelatin coated tissue culture plates pre-seeded with *γ*-irradiated MEF (mouse embryonic fibroblasts), after 2 passages cells were cultivated on 0.1 % gelatin coated tissue culture plates without MEF. Cells were maintained at 37° C, 5% CO_2_, routinely passaged every other day using Trypsin/EDTA detachment, media was replaced every day. Wild-type male embryonic stem cell line v6.5 was purchased from Novus Biologicals (NBP1-41162). Nanog reporter cell line NHET was kindly provided by Jianlong Wang (Icahn School of Medicine, New York, USA). In this cell line, originally generated by Maherali et al. [1] using the design of Hatano et al. [2], the endogenous Nanog open reading frame (ORF) has been substituted by a gene cassette containing green fluorescent protein (GFP) in series with a Puromycin resistance, separated by a internal ribosome entry site (IRES). For 0i and 2i cultures, 3 technical replicates were assessed for Nanog and GFP distributions by flow cytometry and image analysis at passage number 11 (v6.5s) and passage number 20 (NHETs, also day 0 in time-course experiments). For undirected differentiation time-course experiments, three replicates from each initial condition (0i and 2i) were cultured separately for 7 days after withdrawal of LIF from culture media on day 0. Cultures were passaged every 2 days and assessed by flow cytometry and image analysis on days 0, 1, 2, 3, 5 and 7.

### Immunocytochemistry and flow cytometry

Cells for flow cytometry were detached using Trypsin/EDTA. Cells cultures for imaging were briefly washed in PBS. All cells were fixed for 20 minutes at room temperature (RT) in 4% Paraformaldehyde in PBS and washed three times with PBS. Cell and nuclear membranes were permeabilized using 0.1 % Triton-X-100 in PBS for 10 min at RT. Unspecific antibody binding was blocked with 0.1% Triton-X-100 in PBS with 10% fetal bovine serum for 45 min at RT. Blocked cells were washed three times with blocking solution and re-suspended in blocking solution containing either primary antibodies overnight at 4°C. Cell suspensions were under continuous agitation and cell plates were under continuous gentle motion. All experimental results in the main paper used directly conjugated primary antibodies: Mouse anti-mouse Nanog (Alexa Fluor^®^ 647, 1:200, 560279), mouse IgG1*κ* isotype control (Alexa Fluor^®^ 647, 557732), rat anti-histone H3 (pS28) (Alexa Fluor^®^ v450, 560606), rat IgG2a *κ* isotype control, (Alexa Fluor^®^ v450, 560377). Samples were washed three times with PBS and for cell imaging nuclei were incubated with 20*μ*g/ml DAPI (Invitrogen) for 15 mins before imaging. The results shown Fig. S1 also used the following non-conjugated primary antibodies: Mouse anti Oct3/4 (c-10) (Santa Cruz Biotechnology, SC5279), murine IgG2b isotype control (Sigma, SAB4700729). After incubating with primary antibodies overnight these samples were washed 3 times with blocking solution and incubated with a secondary antibodies for 1 hr at RT. Secondary antibodies: goat anti-mouse IgG(H&L) (Alexa Fluor^®^ 488, abcam, abA11017), goat anti-mouse IgG (Alexa Fluor^®^ 647, BioLegend, 405322). Images were recorded using a Nikon Eclipse Ti microscope. Cell suspension samples were analyzed using a BD FACS Aria II fluorescence activated cell sorting device and BD FACS Diva™ software (Becton-Dickinson, Oxford, UK). Flow cytometry analysis was performed using FlowJo^™^, MATLAB^™^ and R [3, 4] software. Nanog and GFP fluorescence was quantified in terms of MESF units (molecules of equivalent soluble fluorophore) using Quantum^™^ Alexa Fluor^®^ 488 and 647 MESF calibration beads (Bangs Laboratories). Fluorescence probability distributions for non-directed differentiation experiments were aligned at the 1st percentile of Nanog and GFP observations between days.

### Image analysis

Image analysis was carried out on grayscale fluorescence images using CellProfiler [5]. Nuclei were identified automatically based on DAPI signal and hand curated to exclude mitotic cells, unresolved or split nuclei and at the image edge. Spatially variable background fluorescence within an image was accounted for using average background subtraction over an image set. Mean fluorescence intensity of Nanog and GFP was calculated for nuclear areas.

### Model fitting

Nanog expression distributions from FACS and image analysis were fitted to a gaussian mixture model with 1 or 2 components using expectation maximization. Model selection was conducted using Bayes Information Criterion (BIC). To ensure that robustly bimodal distributions were identified we required both components to have a weight greater than 0.1 and excluded models in which the peak probability density of one component lay within the other component when fitting to two component mixtures.

### Mutual information calculation

Mutual information (MI) between GFP and Nanog expression levels was estimated using the James-Stein-type shrinkage estimator [6]. The discretization (‘binning’) of each variable at each time point was performed separately via the Bayesian Blocks method [7]. Since MI is invariant to smooth re-parameterization, we worked with the aligned, rescaled log-transformed fluorescence values.

## MATHEMATICAL DETAILS

### Allelic synchronization and mRNA co-expression dynamics

Consider the transcriptional dynamics of 2 alleles of the same gene in a single cell. Let *m*_1_ denote the number of mRNA transcripts associated with allele 1, let *m*_2_ denote the number of mRNA transcripts associated with allele 2, and assume that expression of both alleles are governed by linear birth-death processes with production rates 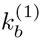, 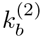 and decay rates 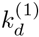, 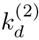. Thus, we are concerned with the dynamics of the following system of reactions:

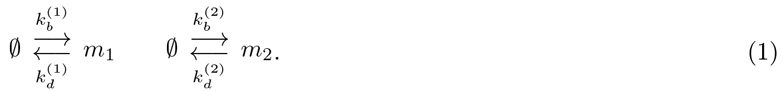

Since the alleles are not coupled together they act independently and the stationary joint probability mass function (PMF) for this process is a prodcut of two independent Poisson processes:

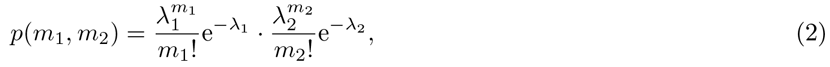

where 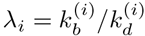. for *i* ∈ {1, 2}. In order to couple the genes together we allow the transcription rates 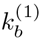 and 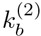 to depend upon the concentration of a shared upstream regulator, gene *X*. Let *x* denote the concentration of *X* and let *ρ*(*x*) be the stationary probability density function for *x*. Taking birth rate as 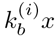, the stationary joint PMF is then obtained from Bayes’ theorem:

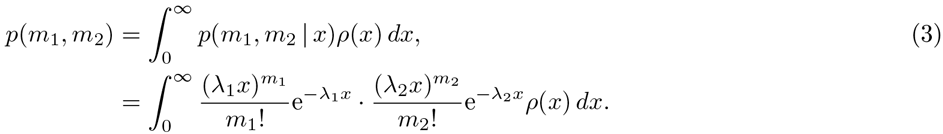

If *x* ~ Gamma(*r*, *θ*) then this gives

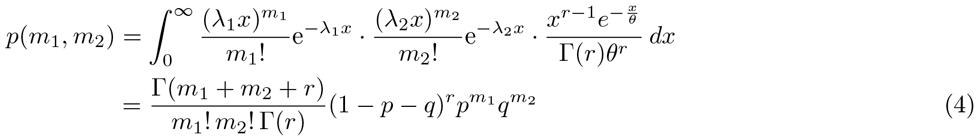

where *p* = λ_1_*θ*/[1 + *θ*(λ_1_ + λ_2_)] and *q* = *ap* with *a* = λ_2_/λ_1_. Thus, the joint PMF is a bivariate negative binomial distribution. Note that the marginal distributions are negative binomial distributions, with probability *p*′ = λ_1_*θ*/(1 + λ_1_*θ*) for allele 1 and *p*″ = λ_2_*θ*/(1 + λ_2_*θ*) for allele 2. For instance, for allele 1:

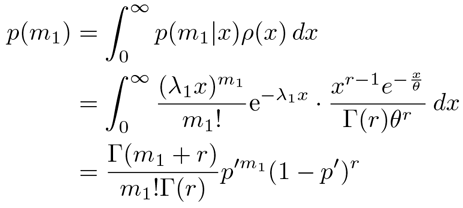

The covariance between *m*_1_ and *m*_2_,

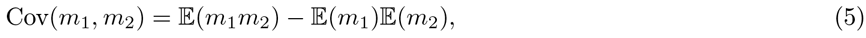

may be obtained from the probability generating function for *p*(*m*_1_, *m*_2_), which in this case is:

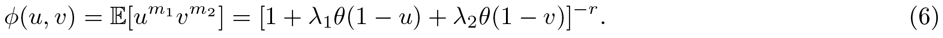

In particular,

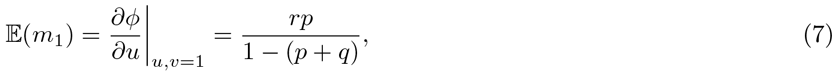

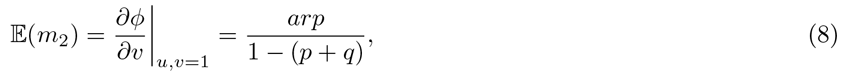

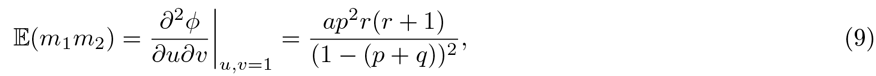

and therefore,

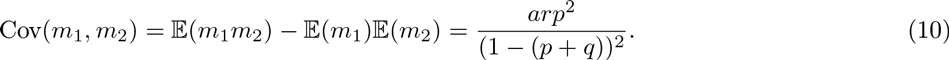

This may be expressed in an alternative form as

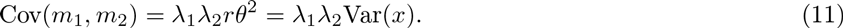

Thus, the covariance of the target genes is proportional to both the variance of the upstream regulator and the sensitivities of the two targets to the upstream regulator. The correlation between *m*_1_ and *m*_2_ may also be similarly calculated. We obtain:

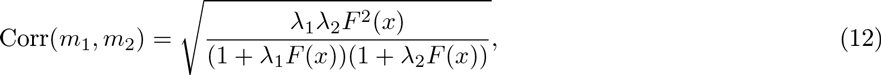

where *F*(*x*) = Var(*x*)/𝔼(*x*) is the Fano factor (also known as the index of dispersion) of the upstream regulator *x*. Since,

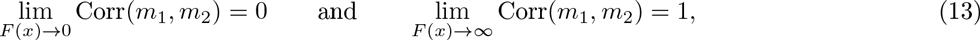

over-dispersion in the upstream regulator increases the correlation between downstream targets and under-dispersion reduces the correlation between targets. If the alleles are kinetically identical (λ_1_ = λ_2_ = λ) then

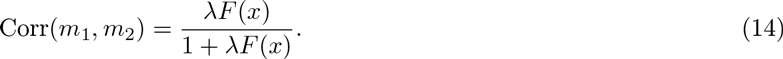

and the correlation between the alleles grows hyperbolically with the dispersion of the upstream regulator.

While the form of joint PMF given in Eq.4 depends upon the upstream regulator being Gamma distributed, Eqs.11–12 hold true for any upstream distribution *ρ*(*x*) with nonnegative support. In general the probability generating function for the joint PMF *p*(*m*_1_, *m*_2_) has the form:

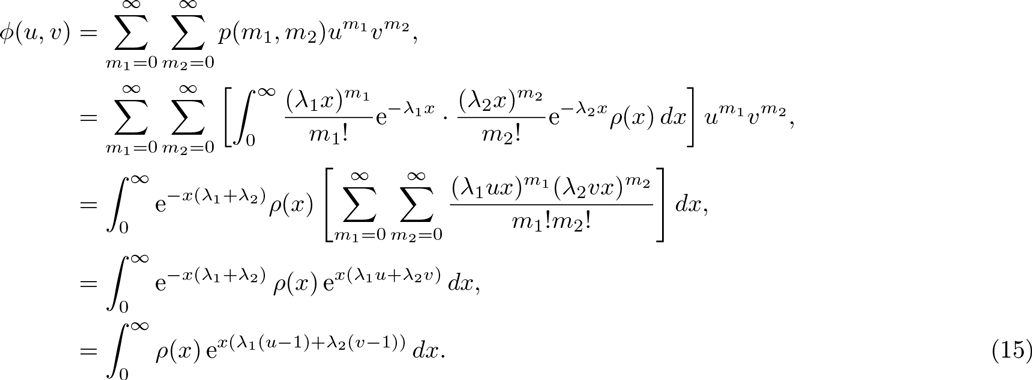

Thus,

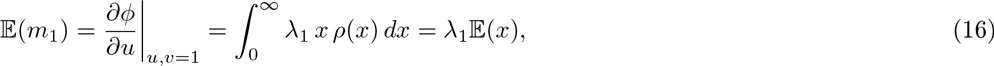

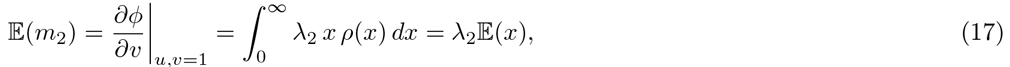

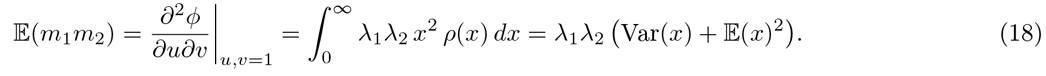

Therefore,

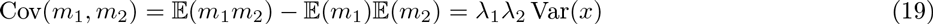

as before. To find the correlation of downstream targets, we also need to find Var(*m*_1_) and Var(*m*_2_). We do so by using:

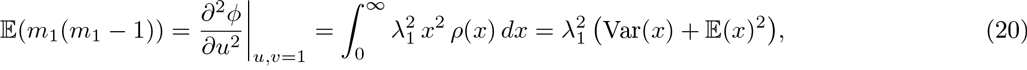

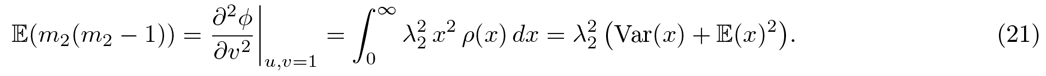

Hence, as 𝔼(*z*(*z* − 1)) = Var(*z*) − 𝔼(*z*) − 𝔼(*z*)^2^ we obtain

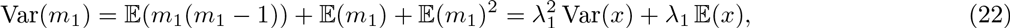

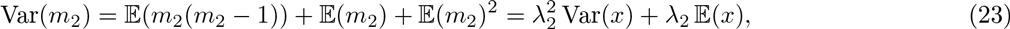

thus giving

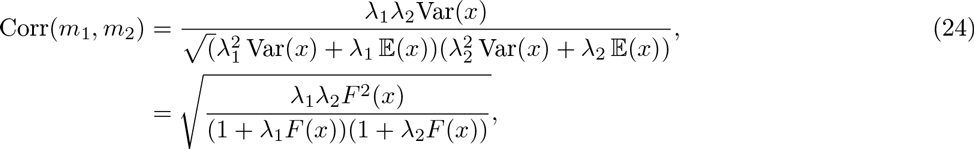

as before.

If the mRNA birth process is not linearly dependent on *x*, but instead is determined by some arbitrary dependence *f*(*x*), then the probability generating function for *p*(*m*_1_, *m*_2_) is given by

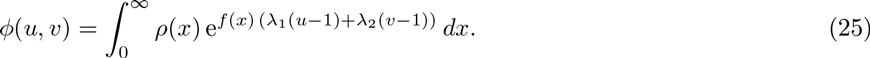

In this case, 𝔼(*m*_1_), 𝔼(*m*_2_), Var(*m*_1_), Var(*m*_2_) and Cov(*m*_1_, *m*_2_) take a similar form as above, but with 𝔼(*f*(*x*)) and Var(*f*(*x*)) replacing 𝔼(*x*) and Var(*x*) respectively. Thus,

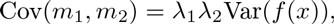

and

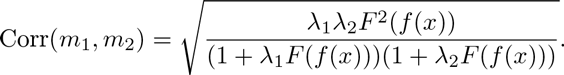

### Bifurcation curves for Nanog dynamics

The dynamics for all the reporter strategies that we consider can be described by the following dimensionless ordinary differential equation (ODE) for total Nanog concentration 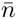 (see main text and below):

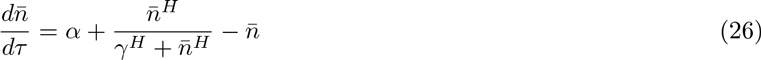

Fixed points solutions, in which 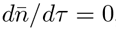, satisfy the polynomial

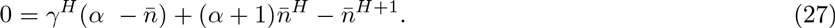

This polynomial has either 1 or three real solutions, depending on the values of *α* and *γ* and the Hill coefficient *H*. When there is only one real solution the system has one stable fixed point and the resulting Nanog is unimodal; when there are three real solutions, two of them are stable and the Nanog distribution is bimodal. The threshold between these regimes occurs when there is a repeated solution, which occurs when the discriminant Δ of Eq.27 is zero. In the case *H* = 2

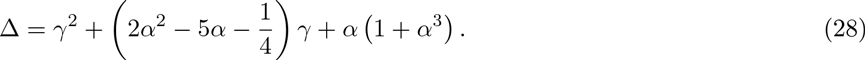

Thus Δ = 0 is a quadratic for *γ* which has roots

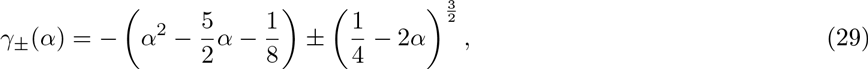

which are the bifurcation curves given in the main text. If the model parameters fall *inside* the region enclosed by these curves then Nanog expression is bimodal, corresponding to the coexistence of a Nanog high and Nanog low expressing subpopulations of cells; if the model parameters fall *outside* this region then Nanog expression is unimodal, corresponding to a homogeneous population of Nanog high or Nanog low expressing cells. A similar calculation may be performed for arbitrary *H* ∈ ℤ^+^.

### Reporter perturbations

To understand how reporters may affect endogeneous Nanog dynamics we compared the dynamics of Nanog in various different reporter lines with those in wild-type ES cells. To recap from the main text, Nanog expression in wild-type cells is described by the following ODEs:

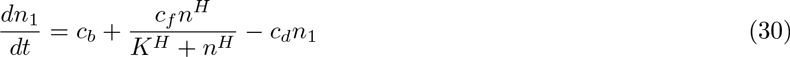

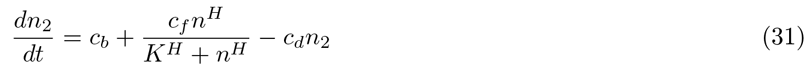

where *n*_*i*_ denotes the concentration of the Nanog protein output of allele *i* ∈ {1, 2}, *n* = (*n*_1_ + *n*_2_) is total Nanog concentration. Combining these equations we obtain an ODE for the total Nanog protein concentration:

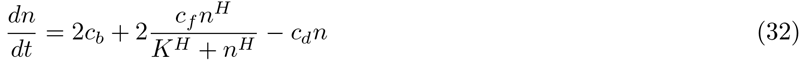

Nondimensionalizing using the scalings 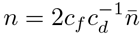, and 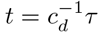 we obtain:

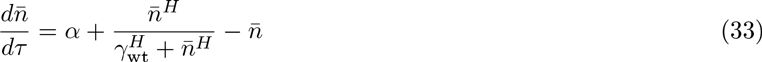

where 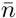 is the dimensionless total Nanog concentration and *τ* is dimensionless time. The dimensionless constants *α* = *c*_*b*_/*c*_*f*_ and *γ* = *γ*_wt_ = *c*_*d*_*K*/2*c*_*f*_ measure the strength of the baseline production rate and the strength of the Nanog autoregulatory feedback loop respectively. We now consider how Nanog dynamics given by Eq.33 are perturbed by a variety of different kinds of reporters. In all cases, for clarity of exposition, the reporter proteins are assumed to decay with the same kinetics as Nanog. This assumption may be weakened without affecting our conclusions. The results of this section are also summarised in Supplementary Tables 1 & 2 which also give the relationships between reporter and Nanog concentrations at equilibrium as a measure of the quantitative accuracy of each reporter.

### Single Allele Reporter Strategies

#### Knock-in reporters

Knock-in reporters reporters remove the Nanog protein coding region from one allele and replace it with a reporter gene under the same promoter control. In this case, the kinetics described by Eqs.33 are modified to:

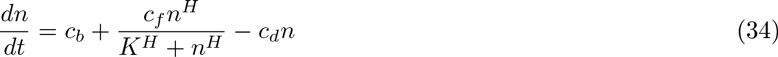

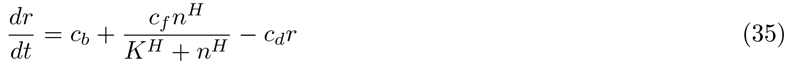

where *n* denotes Nanog concentration and *r* denotes reporter concentration. Using the scalings 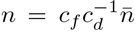 and 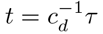 the dimensionless equation for total Nanog concentration in the knock-in reporter line is:

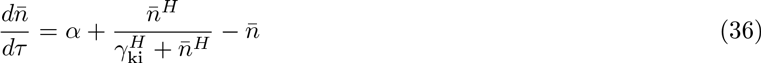

where *α* is as before, but *γ*_ki_ = 2*γ*_wt_. In this case, the loss of Nanog production from one allele has the effect of diminishing the functional Nanog production rate by a factor of two, which effectively weakens the endogenous feedback mechanisms and thereby doubles *γ*. Since the magnitude of *γ* determines if Nanog is homogeneously or heterogeneously expressed in the population, this change can induce a heterogeneous Nanog expression pattern in a reporter cell line that is not characteristic of the wild-type (or vice versa). From Eqs.34–35 at equilibrium *r* = *n* so it is expected that the knock-in reporter signal will faithfully represent Nanog expression in the engineered line.

#### Pre/post (PP) reporters

Single allele pre/post reporters insert the reporter gene either directly before or after the Nanog protein coding region on one Nanog allele. We assume that this insertion reduces the Nanog production rate from the reporter allele by a factor 0 ≤ *ϵ* ≤ 1 due to increased transcription time. The following ODEs describe the dynamics in this case:

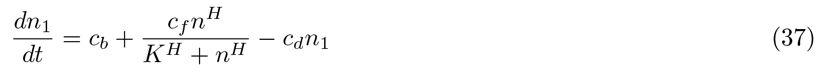

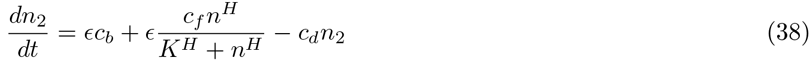

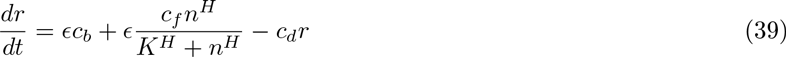

where *n_i_* denotes the concentration of the Nanog protein output of allele *i* ∈ {1, 2} and *r* denotes the reporter concentration (assumed without loss of generality to be produced from allele 2). Combining these equations for total Nanog *n* = *n*_1_ + *n*_2_ and using the scalings 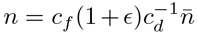 and 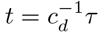, the dimensionless equation for total Nanog is:

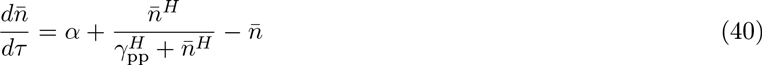

where *γ*_pp_ = 2*γ*_wt_/(1 + *ϵ*). Thus, if the addition of the reporter gene completely halts transcription from allele 2 then *ϵ* = 0 and *γ*_pp_ = *γ*_ki_ = 2*γ*_wt_; if the reporter halves the rate of transcription from allele 2 then *ϵ* = 1/2 and *γ*_pp_ = 4*γ*_wt_/3; if the reporter does not affect the rate of transcription from allele 1, then *ϵ* = 1 and *γ*_pp_ = *γ*_wt_. For 0 < *ϵ* < 1 pre/post reporters are less likely than knock-in reporters to induce qualitative changes in Nanog expression dynamics, yet are still subject to similar systemic risk. From Eqs.37–39 at equilibrium *r* = *ϵn*/(1 + *ϵ*) so it is expected that, in addition to any qualitative perturbations, the PP reporter signal will quantitatively misrepresent Nanog expression by a factor *ϵ*/(1 + *ϵ*).

#### Multiple pre/post (MPP) reporters

If multiple (*m*) repeats of the reporter gene are inserted on the reporter allele then production rates are reduced by a factor of *ϵ*_*m*_ where 0 ≤ *ϵ*_*m*_ ≤ ϵ ≤ 1 and *m* copies of the reporter transcript are produced (for non-tandem repeats). The dynamics become

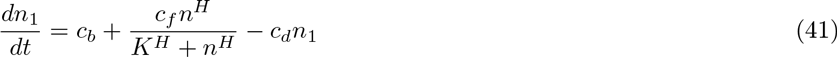

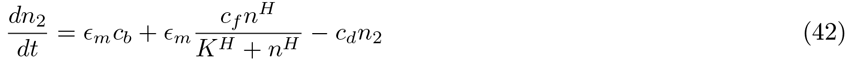

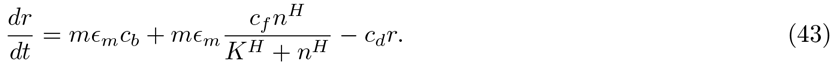

Combining these equations and using the scalings 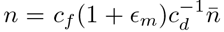 and 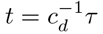, the dimensionless equation for total Nanog is:

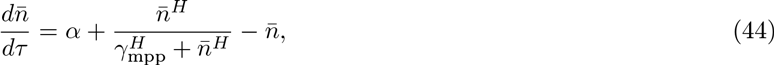

where *γ*_mpp_ = 2*γ*_wt_/(1 + *ϵ*_*m*_). If a single insert slows transcription by a factor 0 ≤ *ϵ* ≤ 1 (as with the PP reporter) and each of the *m* inserts is identical, then *ϵ*_*m*_ = *ϵ*/(*m*(1 − *ϵ*) + *ϵ*) ≤ *ϵ* (with equality if and only if *m* = 1). Thus, although multiple reporter additions improve fluorescent signal, the systemic risk is increased with each additional reporter insert. As *m* becomes large *ϵ*_*m*_ → 0 and this risk approaches that of the knock-in reporters. From Eqs.41–43 at equilibrium *r* = *mϵ_m_n*/(1 + *ϵ*_*m*_) so it is expected that, in addition to any qualitative perturbations, the MPP reporter signal will quantitatively misrepresent Nanog expression by a factor *mϵ_m_*/(1 + *ϵ*_*m*_).

#### Fusion reporters

Fusion reporters produce a modified version of Nanog, which includes a fluorescence structure as part of the Nanog protein. For single allele fusion reporters, the fusion protein (concentration *n*_2_) is produced from one allele, and the wild-type protein (concentration *n*_1_) is produced from the other. This has two effects on the dynamics: (1) the rate of transcription from the reporter allele is reduced by a factor 0 ≤ *ϵ* ≤ 1 due to the additional DNA that must be transcribed, as for a PP reporter, and (2) the function of the Nanog from the reporter allele is compromised by a factor 0 < *δ* < 1 due to the addition of a cumbersome fluorescent protein to the native Nanog. The dynamics in this case are:

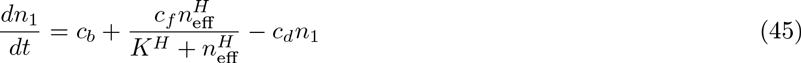

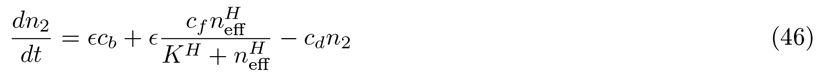

where *n*_eff_ = *n*_1_ + *δn_2_*. Combining these equations and nondimensionalising using the scalings 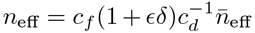 and 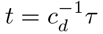 we obtain the following equation for the effective Nanog concentration:

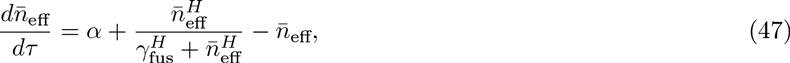

where *γ*_fus_ = 2*γ*_wt_/(1 + *ϵδ*). If *δ* = 0 then the Nanog-reporter fusion is not functional, while for *δ* = 1 the Nanog-reporter fusion functions as the native Nanog protein. For 0 < *δ* < 1, *γ*_fus_ > *γ*_pp_, therefore fusion reporters are more likely than pre/post reporters to induce qualitative changes in expression dynamics. From Eqs.45-46 at equilibrium *r* = *ϵn*/(1 + ϵ) so it is expected that, in addition to any qualitative perturbations, the fusion reporter signal will quantitatively misrepresent Nanog expression by a factor *r* = *ϵ*/(1 + *ϵ*).

#### BAC reporters

Bacterial artificial chromosome (BAC) reporters introduce a piece of extra-genomic DNA into the cell that encodes the Nanog gene under the control of the endogenous Nanog promoter and regulatory regions. Because this construct does not disturb the kinetics of either of the wild-type alleles, it (uniquely amongst the reporters we consider) does not directly affect the endogenous feedback mechanisms and is therefore the least likely reporter strategy to induce qualitative changes in Nanog dynamics. However, because the reporter construct is physically separated from the Nanog alleles, it is expected that the reporter protein expression is subject to extrinsic stochastic fluctuations which are independent to those of endogenous Nanog expression. For this reason we expect that BAC reporters are more susceptible to *technical* errors that the other constructs we consider.

### Dual Allele Reporter Strategies

Dual allele reporters can either express the same reporter molecule from both allele (e.g. both drive transcription of GFP) or may express different reporter molecules from different alleles (e.g. GFP from one allele, and a red fluorescent protein from the other). The analysis of the single allele reporters above may be easily modified to account for dual reporter strategies. The dynamics for total Nanog in dual reporter systems are given by:

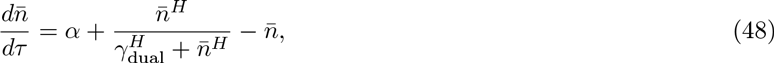

where: (1) *γ*_dual_ = *γ*_wt_/*ϵ*_*m*_ in the case of dual multiple pre/post reporters that produce m copies of the same fluorescent signal from both alleles (which reduces the rate of transcription from both alleles by a factor 0 ≤ *ϵ*_*m*_ ≤ 1 from both alleles); (2) *γ*_dual_ = 2*γ*_wt_/(*ϵ*_*m*1_ + *ϵ*_*m*2_) in the case of dual pre/post reporters that produce different reporters from the two alleles (which reduces the rate of transcription from alleles 1 and 2 by factors 0 < *ϵ*_*m*1_ ≤ 1 and 0 ≤ *ϵ*_*m*2_ ≤ 1 respectively); (3) *γ*_dual_ = *γ*_wt_/*ϵδ* for dual fusion reporters that produce the same fusion protein from each allele (which reduces the rate of transcription from both alleles by a factor 0 ≤ *ϵ* ≤ 1 from both alleles, and compromises Nanog function by a factor *δ*); (4) *γ*_dual_ = 2*γ*_wt_/(*ϵ*_1_*δ*_1_ + *ϵ*_2_*δ*_2_) for dual fusion reporters that produce different fusion proteins from each allele (which reduce the rates of transcription by factors 0 ≤ *ϵ*_1_ ≤ 1 and 0 ≤ *ϵ*_2_ ≤ 1 from alleles 1 and 2 respectively, and compromise Nanog function by factors *δ*_1_ and *δ*_2_ respectively). It should be noted for dual allele fusion reporters there is no wild-type protein in the system at all which could have further unintended consequences, including off target effects. In all cases *γ*_dual_ is larger than the corresponding value of *γ* for the single allele reporters. Thus, while more technically accurate, dual allele reporters carry raised risk of systemic perturbations to the endogenous kinetics. In addition to any qualitative perturbations, dual reporter systems may also quantitatively misrepresent Nanog expression in similar ways to the corresponding single allele constructs. These perturbations are detailed in Supplementary Table 2.

## SUPPLEMENTARY FIGURE AND TABLE CAPTIONS

### Supplementary Figure 1

**Nanog and Oct3/4 immunofluorescence. A** Grayscale and composite RGB images of DAPI staining, Nanog immunofluorescence and GFP fluorescence in v6.5 and NHET cells in 0i and 2i conditions. Boxes indicate the regions of the image shown in Figure 1 main text. Variability of Nanog fluorescence can be seen for both v6.5 and NHET cells, with substantially greater variability in 0i conditions. **B** Grayscale and composite RGB images for Oct3/4 immunofluorescence from v6.5 and NHET cells in 0i and 2i cultures. Oct3/4 is less variably expressed than Nanog. **C** Grayscale and composite RGB images for Nanog and Oct3/4 antibody isotype controls for v6.5 0i cells.

### Supplementary Figure 2

**Image analysis of Nanog variability in v6.5 and NHET cells. A** Examples of Nanog immunofluorescence distributions for v6.5 cells and joint Nanog-GFP distributions for NHET cells in 0i and 2i conditions, assessed by image analysis. NHET populations are split by GFP expression (high/low) and Nanog expression (highest 20% /lowest 20%). Percentages are shown for outer subpopulations. **B**. Assessment of joint Nanog-GFP distributions in NHET cells during differentiation subsequent to LIF-withdrawal starting from 0i and 2i cultures, using data assessed by image analysis. **C**. Mutual information between Nanog and GFP during differentiation, using data assessed by image analysis.

### Supplementary Table 1

**Perturbations by single allele reporters.** Summary of analysis of reporter perturbations to wild-type dynamics for single allele reporters.

### Supplementary Table 2

**Perturbations by dual allele reporters.** Summary of analysis of reporter perturbations to wild-type dynamics for dual allele reporters.

## References

[1] Prasher DC, Eckenrode VK, Ward WW, Prendergast FG, Cormier MJ (1992) Primary structure of the Aequorea victoria green-fluorescent protein. Gene 111(2):229–233.

[2] Chalfie M, Tu Y, Euskirchen G, Ward WW, Prasher DC (1994) Green fluorescent protein as a marker for gene expression. Science 263(5148):802–805.

[3] Chalfie M, Kain SR (2005) Green Fluorescent Protein: Properties, Applications and Protocols. (John Wiley and Sons, New Jersey), Second edition.

[4] Faddah DA et al. (2013) Single-cell analysis reveals that expression of nanog is biallelic and equally variable as that of other pluripotency factors in mouse ESCs. Cell Stem Cell 13(1):23–9.

[5] Filipczyk A et al. (2013) Biallelic expression of Nanog protein in mouse embryonic stem cells. Cell Stem Cell 13(1):12–3.

[6] Wigner E (1963) The problem of measurement. American Journal of Physics 31(1):6–15.

[7] Chambers I et al. (2007) Nanog safeguards pluripotency and mediates germline development. Nature 450(7173):1230–4.

[8] Miyanari Y, Torres-Padilla ME (2012) Control of ground-state pluripotency by allelic regulation of Nanog. Nature 483(7390):470–3.

[9] Abranches E, Bekman E, Henrique D (2013) Generation and characterization of a novel mouse embryonic stem cell line with a dynamic reporter of Nanog expression. PloS one 8(3):e59928.

[10] Abranches E et al. (2014) Stochastic NANOG fluctuations allow mouse embryonic stem cells to explore pluripotency. Development (Cambridge, England) 141(14):2770–2779.

[11] Friedman N, Cai L, Xie XS (2006) Linking stochastic dynamics to population distribution: An analytical framework of gene expression. Physical Review Letters 97(Oct):1–4.

[12] Kalmar T et al. (2009) Regulated fluctuations in nanog expression mediate cell fate decisions in embryonic stem cells. PLoS biology 7(7):e1000149.

[13] Canham MA, Sharov AA, Ko MSH, Brickman JM (2010) Functional heterogeneity of embryonic stem cells revealed through translational amplification of an early endodermal transcript. PLoS Biology 8(5):e1000379.

[14] Martinez Arias A, Brickman JM (2011) Gene expression heterogeneities in embryonic stem cell populations: Origin and function. Current Opinion in Cell Biology 23(6):650–656.

[15] Cahan P, Daley GQ (2013) Origins and implications of pluripotent stem cell variability and heterogeneity. Nature Reviews. Molecular Cell Biology 14(6):357–68.

[16] Torres-Padilla ME, Chambers I (2014) Transcription factor heterogeneity in pluripotent stem cells: a stochastic advantage. Development 141(11):2173–2181.

[17] Wang J et al. (2006) A protein interaction network for pluripotency of embryonic stem cells. Nature 444(7117):364–368.

[18] Kim J, Chu J, Shen X, Wang J, Orkin SH (2008) An extended transcriptional network for pluripotency of embryonic stem cells. Cell 132(6):1049–1061.

[19] MacArthur BD, Ma’ayan A, Lemischka IR (2009) Systems biology of stem cell fate and cellular reprogramming. Nature Reviews. Molecular Cell Biology 10(10):672–681.

[20] Dunn SJ, Martello G, Yordanov B, Emmott S, Smith AG (2014) Defining an essential transcription factor program for naïve pluripotency. Science 344(6188):1156–1160.

[21] MacArthur BD, Please CP, Oreffo ROC (2008) Stochasticity and the molecular mechanisms of induced pluripotency. PloS One 3(8):e3086.

[22] Glauche I, Herberg M, Roeder I (2010) Nanog variability and pluripotency regulation of embryonic stem cells-insights from a mathematical model analysis. PloS One 5(6):e11238.

[23] Wang J, Levasseur DN, Orkin SH (2008) Requirement of Nanog dimerization for stem cell self-renewal and pluripotency. Proceedings of the National Academy of Sciences of the United States of America 105(17):6326–6331.

[24] MacArthur BD et al. (2012) Nanog-dependent feedback loops regulate murine embryonic stem cell heterogeneity. Nature Cell Biology 14(11):1139–1147.

[25] Xiong W, Ferrell JEJ (2003) A positive-feedback-based bistable ‘memory module’ that governs a cell fate decision. Nature 426(6965):460–465.

[26] Tyson JJ, Chen KC, Novak B (2003) Sniffers, buzzers, toggles and blinkers: Dynamics of regulatory and signaling pathways in the cell. Current Opinion in Cell Biology 15:221–231.

[27] Becskei A, Seraphin B, Serrano L (2001) Positive feedback in eukariotic gene networks: Cell differentiation by graded to binary response conversion. The EMBO Journal 20(10):2528–2535.

[28] Silva J et al. (2009) Nanog is the gateway to the pluripotent ground state. Cell 138(4):722–737.

[29] Maherali N et al. (2007) Directly Reprogrammed Fibroblasts Show Global Epigenetic Remodeling and Widespread Tissue Contribution. Cell Stem Cell 1(July):55–70.

[30] Ying QL et al. (2008) The ground state of embryonic stem cell self-renewal. Nature 453(7194):519–523.

[31] Cover TM, Thomas JA (2006) Elements of Information Theory. (John Wiley and Sons, New Jersey), Second edition.

[32] Snapp E (2005) Design and use of fluorescent fusion proteins in cell biology. Current Protocols in Cell Biology chapter 21, Unit 21.4.

[33] Day RN, Davidson MW (2009) The Fluorescent Protein Palette: Tools for Cellular Imaging. Chemical Society Reviews 38(10):2887–2921.

## References

[1] Maherali N et al. (2007) Directly Reprogrammed Fibroblasts Show Global Epigenetic Remodeling and Widespread Tissue Contribution. Cell Stem Cell 1(July):55–70.

[2] Hatano SY et al. (2005) Pluripotential competence of cells associated with Nanog activity. Mechanisms of development 122(1):67–79.

[3] R Core Team (2016) R: A Language and Environment for Statistical Computing (R Foundation for Statistical Computing, Vienna, Austria).

[4] Wickham H (2009) ggplot2: Elegant Graphics for Data Analysis. (Springer-Verlag New York).

[5] L K et al. (2011) Improved structure, function, and compatibility for cellprofiler: modular high-throughput image analysis software. Bioinformatics 27:1179–1180.

[6] Hausser J, Strimmer K (2009) Entropy inference and the James-Stein estimator, with application to nonlinear gene association networks. Journal of Machine Learning Research 10(June):1469–1484.

[7] Scargle JD (1998) Studies in Astronomical Time Series Analysis. V. Bayesian Blocks, a New Method to Analyze Structure in Photon Counting Data. The Astrophysical Journal 504(1):405–418.

